## Supporting information for "Differentiation of leukocytes in bronchoalveolar lavage fluid samples using higher harmonic generation microscopy and deep learning"

**S1 Table: Evaluation of top-performing ResNet50 model on the validation set (top rows, light grey) and testing set (bottom rows, dark grey).** The average mean absolute error on the validation set was 0.136. The average mean absolute error on the testing set was 0.087. For each case in the two sets, the standard cytology percentages with standard deviation (except for the PBMC in the testing set, because this sample was counted once) are given and compared to the model’s output. Given that a mosaic may be divided into multiple, single cases, an average regression output and standard deviation can be derived. For the blood fraction data, the macrophages class is replaced by monocytes, since monocytes are the precursor cells of macrophages present in blood.

| **Sample** | **Regression** | **Neutrophils (%)** | **Eosinophils (%)** | **Lymphocytes (%)** | **Monocytes / Macrophages (%)** |
| --- | --- | --- | --- | --- | --- |
| **GR 2** | **Cytology** | 94.0 ± 1.3 | 5.0 ± 0.5 | 1.0 ± 0.8 | 0.0 ± 0.0 |
|  | **ResNet50** | 52.7 ± 3.2 | 20.9 ± 3.2 | 15.0 ± 1.6 | 11.6 ± 1.7 |
| **PBMC 2** | **Cytology** | 1.9 ± 0.8 | 0.0 ± 0.0 | 94.6 ± 0.6 | 3.5 ± 1.3 |
|  | **ResNet50** | 6.4 ± 1.3 | 0.8 ± 0.3 | 80.6 ± 3.1 | 12.3 ± 1.6 |
| **BALF 2** | **Cytology** | 17.6 ± 2.3 | 4.2 ± 0.1 | 4.7 ± 0.8 | 73.5 ± 2.9 |
|  | **ResNet50** | 6.7 ± 1.1 | 0.4 ± 0.1 | 11.6 ± 1.3 | 81.4 ± 1.3 |
| **GR 3** | **Cytology** | 88.3 ± 1.9 | 10.6 ± 1.1 | 1.1 ± 0.8 | 0.0 ± 0.0 |
|  | **ResNet50** | 77.0 ± 5.5 | 6.7 ± 2.7 | 6.0 ± 1.5 | 9.3 ± 3.1 |
| **PBMC 3** | **Cytology** | 2.9 | 0.6 | 95.3 | 1.2 |
|  | **ResNet50** | 27.4 ± 10.6 | 1.4 ± 0.8 | 61.7 ± 11.4 | 9.5 ± 1.5 |
| **BALF 4** | **Cytology** | 32.0 ± 0.9 | 4.2 ± 0.5 | 6.4 ± 0.6 | 57.4 ± 1.0 |
|  | **ResNet50** | 22.5 ± 2.9 | 2.6 ± 0.6 | 9.5 ± 2.4 | 65.5 ± 2.4 |
